# Enhanced Prediction of Seafloor Ecological State Using 16S Nanopore Sequencing

**DOI:** 10.1101/2024.10.25.620171

**Authors:** Melcy Philip, Tonje Nilsen, Sanna Majaneva, Ragnhild Pettersen, Morten Stokkan, Jessica Louise Ray, Nigel Keeley, Knut Rudi, Lars-Gustav Snipen

## Abstract

Anthropogenic stress on benthic habitats, particularly from aquaculture, calls for accurate and efficient monitoring of the macrofauna ecological state. Recent advancements in Oxford Nanopore Technology (ONT) together with environmental DNA offers cost-effective and rapid, on-site monitoring of such ecosystems. Previous studies have demonstrated that Nanopore sequencing provides sufficient precision for predicting ecological state, despite reported challenges with sequencing accuracy. In this study, we aim to predict the seafloor ecological state with both Illumina and Nanopore 16S rRNA gene sequencing data and using a combination of machine learning and feature selection. We analyzed 88 seafloor samples from aquaculture sites located on a north-south gradient along the Norwegian coast. Both sequencing methods were evaluated in combination with various bioinformatic approaches in the context of predicting the normalized EQR index (nEQR, standard ecological index based on macroinvertebrate counting) as a metric of seafloor ecosystem status. Our results show that the predictive performance of Illumina and Nanopore sequencing platforms are comparable, establishing Nanopore as a feasible alternative to illumina sequencing. By employing a stabilized LASSO regression, the feature set (potential taxa) was efficiently optimized from thousands to 40-60 OTUs. The feature selection reduced prediction errors to less than half of what was obtained through full feature modeling. This feature set demonstrated strong predictive accuracy across both sequencing technologies, with a high correlation between observed and predicted nEQR values. The Pearson correlation coefficient of 0.98 for Illumina and 0.95 (mean prediction error: ±0.04) for Nanopore data (mean prediction error: ±0.06). This study demonstrates that continual improvements in Nanopore sequencing accuracy, in combination with optimized feature selection on a broader set of samples, provides a precise and cost-effective monitoring method for marine benthic environments.

## Introduction

Coastal marine benthic ecosystems serve a central role in biogeochemical cycling and ecosystem services, with pivotal roles in nutrient cycling, sediment dynamics, and maintaining the equilibrium of marine environment (Mäkelin et al., 2024; Montserrat et al., 2008). Increasing frequency and intensity of anthropogenic stressors have led to various shifts in benthic composition and biodiversity (Bauer et al., 2013; Halpern et al., 2015; Lin et al., 2024). The discharge of excess fish feed and waste together with other organic materials from fin fish aquaculture installations have been identified as a significant factor contributing to the irreversible or reversible destruction of benthic ecosystems (González-Gaya et al., 2022; Wang & Olsen, 2024). The rapid changes observed in marine ecosystems call for frequent and efficient surveys to monitor the health of impacted benthic environments to allow early mitigation (González-Gaya et al., 2022; Smit et al., 2021).

From the various abiotic and biotic indicators existing for marine benthic health monitoring, ecosystem indexing based on the abundance of benthic macroinvertebrates is a widely recognised strategy (Borja, 2004; Borja et al., 2004; Karr, 1981; Ruaro & Gubiani, 2013). The normalised Ecological Quality Ratio (nEQR) values derived from morphology-based characterization of benthic communities provide a metric for scoring the ecological status of macrofaunal communities. The nEQR index is specifically calibrated for Norway, tailored to Norwegian regions and water types (Vanndirektivet, 2009). The indexing follows Norwegian guidelines, rather than utilizing standard AMBI index of Borja (Hess et al., 2020). The nEQR index is for practical reasons often grouped into five discrete categories: very poor (0 - 0.2), poor (0.2 - 0.4), moderate (0.4 - 0.6), good (0.6 - 0.8) and very good (0.8 - 1.0). Morphology-based characterization analysis is, however, costly, time-consuming, and require special taxonomic knowledge, prompting the exploration of more efficient monitoring approaches that complement current methods. This includes environmental DNA (eDNA) based approaches which employs short and long read technologies to generate prokaryote and/or eukaryote community profiles of investigated samples (Bohmann et al., 2014; Elahi et al., 2015; TABERLET et al., 2012; Zorz et al., 2023).

Benthic microbial communities are sensitive indicators of ecosystem health, exhibiting distinct patterns in response to environmental, biological, and anthropogenic stressors (Bonthond et al., 2023; Kerrigan & D’Hondt, 2022; Orsi, 2018). Studies to assess the impact of aquaculture on the benthos observed that patterns in the microbial composition correspond to patterns of macrofaunal abundance in the area, the latter of which represent the current gold standard for assessment of ecological condition (Dowle et al., 2015; Pettersen et al., 2022). Advancements in long-read sequencing technologies, such as Oxford Nanopore Technology (ONT), have enabled the generation of more comprehensive and high-resolution microbial profiles (Kerkhof, 2021). Simultaneously, studies that have combined Illumina sequencing technology with machine learning, have demonstrated enhanced prediction of ecosystem health by efficiently processing complex microbial profiles and linking them to key ecological indicators, such as macrofauna abundance (Cordier et al., 2017; Frühe et al., 2021; Wijaya et al., 2023; Wilhelm et al., 2022). Thus, we hypothesize that similar integration of long-read Nanopore technology with machine learning can also facilitate enhanced prediction of ecosystem state, despite its error rates (Wang et al., 2021), enabling more frequent monitoring.

In this study, we compare Nanopore and Illumina technologies to evaluate their effectiveness in generating eDNA signals for predicting normalized Ecological Quality Ratios (nEQR) macrofauna index as the gold standard measure of ecosystem health in Norway. Different methods for obtaining microbial community abundance profiles from both sequence data types are assessed, alongside machine learning strategies for predicting the nEQR index (Figure 1). The work also implements feature selection on microbial read count matrices with the aim to discover microorganisms that are consistently selected across many algorithms and models (Figure 1).

**Figure 1:**
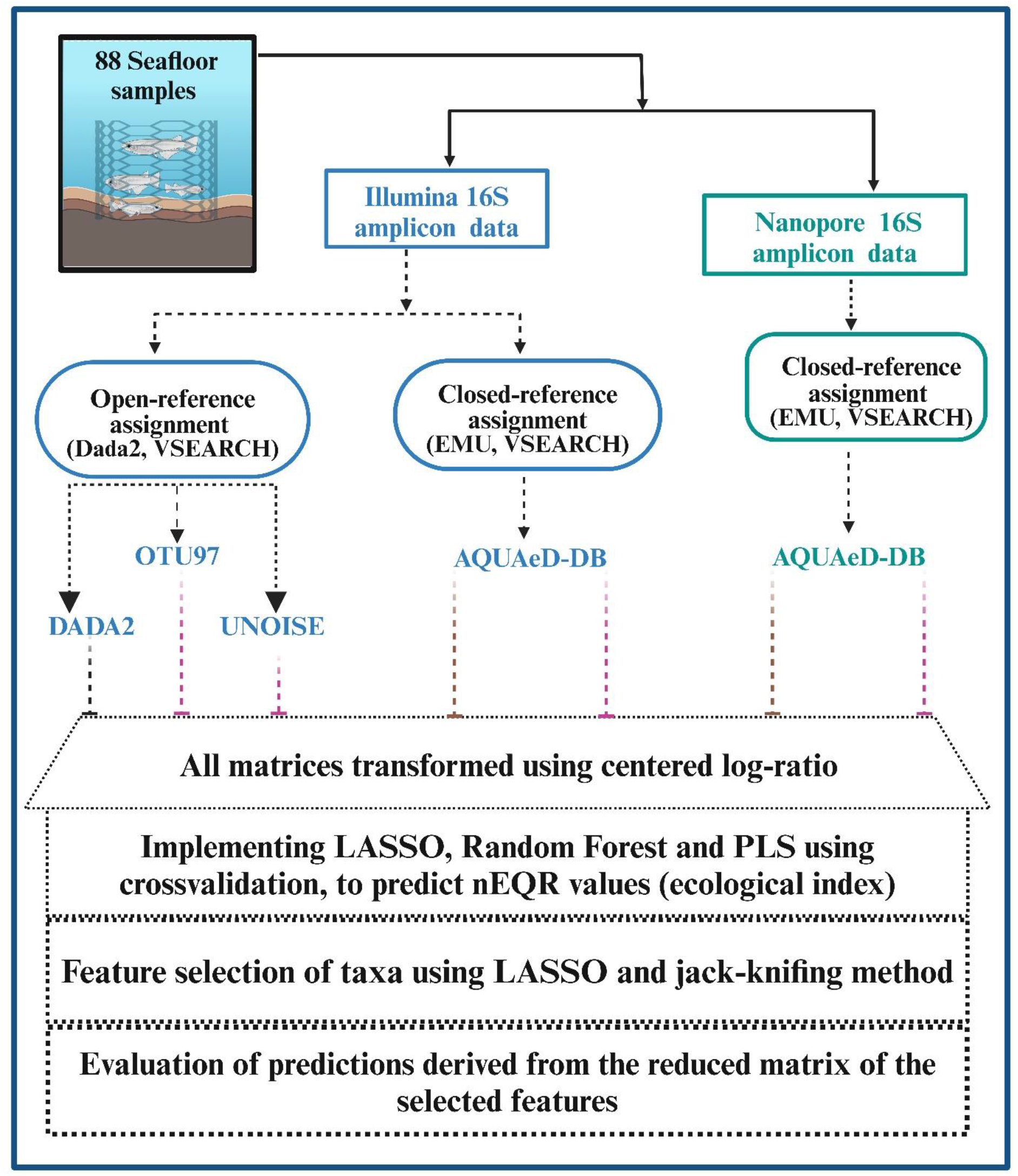
Illustration of bioinformatic workflow. Eighty-eight seafloor samples were sequenced using Illumina and Nanopore technologies. Illumina sequences were assigned using both open- and closed-reference methods, while Nanopore sequences were assigned using closed-reference methods only. Closed-reference assignment methods utilized the AQUAeD-DB database (Philip et al., 2024) as reference for taxonomic assignment of OTUs. Readcount matrices were centered log-ratio transformed prior to predictive modelling (LASSO, Random Forest, PLS) with cross-validation. Feature selection was conducted using LASSO and jack-knifing, followed by prediction evaluation based on selected features.

## Materials and Methods

### Data

A total of 88 seafloor sediment samples were collected from 25 fish farm localities along the Norwegian coast in conjunction with the [AQUAeD project](https://www.nmbu.no/forskning/prosjekter/aquaed). These samples were subjected to 16S amplicon sequencing using Illumina short-read technology targeting the V3-V4 region (∼430 bases) as well as Oxford Nanopore long-read technology targeting the V3-V9 region (∼1100 bases). The sampling, DNA extraction and sequencing protocols are described in detail in a previous publication (Philip et al., 2024). Sequence data were split for cross-validation (see Section e-DNA profiling) based on the locality information provided in the metadata, including the most recent nEQR score for each site.

### e-DNA profiling

Illumina read were assigned using both closed-reference (assignment of reads to a reference database) and open-reference (the sequences are clustered de novo) assignment methods (Fig. 1) (Rideout et al., 2014). The closed-reference approach was used for Nanopore data (Fig. 1) as this method accommodates the higher sequencing error rates commonly observed for Nanopore sequence data. The Nanopore reads from the same samples were filtered by length retaining reads of length 800 - 1400 bases.

The closed-reference approach for Illumina and Nanopore analyses were performed using the VSEARCH software (v2.28.1) (Rognes et al., 2016) and the EMU software (Curry et al., 2022) with the AQUAeD-DB (Philip et al., 2024), consisting of 14,545 reference sequences, obtained from seafloor communities along the Norwegian coast. Reads from both sequencing technologies (Illumina and Nanopore) were assigned to the reference database using both tools (VSEARCH and EMU). This resulted in four read count matrices for each sample.

The open-reference assignment of Illumina sequences was also included since this is the most common approach when using Illumina data. It was performed using two different software tools: DADA2 (v1.28.0) (Callahan et al., 2016) and VSEARCH, the latter with both identity clustering (97%) and denoising using the UNOISE algorithm (Edgar, 2016). This resulted in the ‘Dada2’ read count matrix for Amplicon Sequence Variants (ASVs), the ‘OTU97’ matrix based on 97% identity clustering and the ‘UNOISE’ matrix. All matrices were transformed using Aitchison centered log-ratio (Aitchison, 1982), using 1 pseudo read count. This is common practice when using such compositional data for machine learning (Huang et al., 2023).

### Machine learning approaches

Three standard Machine Learning (ML) algorithms were implemented on all datasets: (1) Least Absolute Shrinkage and Selection Operator (LASSO) (Tibshirani, 1996) from the glmnet package in R, (2) Random Forest (RF) (Segal, 2004) from the randomForest package in R and (3) Partial Least Square Regression (PLSR) (Wold, 1966) from the pls package in R (Liland et al., 2024). Cross-validation by locality was performed for all three approaches. Prediction errors for nEQR values, calculated as the absolute difference between predicted and observed values, were evaluated to assess the impact of various technologies and tools. Additionally, an analysis of variance (ANOVA) was conducted to assess the significant effects on the prediction errors.

### Feature selection

Readcount matrices from community profiles generally exhibit high dimensionality, with many features representing taxa whose abundance may or may not contribute toward observed changes in nEQR indices, and thus, environmental status. Feature selection, in this context, aims to identify the taxa whose abundance makes the largest contribution to nEQR prediction. However, this is known to be an inherently unstable problem, where small variations in the data can lead to different sets of selected features (Hernández Medina et al., 2022). To address this instability, we employed a jackknife resampling approach (Taylor & Kim, 2011). In this approach, the samples were divided by 25 distinct locations, and during each iteration, the samples from one location were systematically left out. Feature selection was then performed on the remaining samples using the LASSO method, resulting in 25 different sets of selected features. The frequency with which each feature was selected across the 25 iterations was recorded, ranging from 0 to 25. Features that were selected frequently across iterations are more likely to represent biologically meaningful taxa, as their selection remained consistent despite slight variations in the data across iterations. The features that appeared at least 50% of the times across all folds (13 out of 25) were selected for further analysis. The read counts for these selected features were then extracted and used for machine learning. The prediction errors were subsequently assessed.

The predictive power of selected features as a function of sequencing technology (Illumina or Nanopore) was also assessed. Correlation between observed and predicted nEQR values was evaluated in two scenarios: First, features selected from a data set were extracted from their respective datasets and used for prediction. Second features selected from Illumina data were extracted from the Nanopore data, and vice versa, in order to see if their predictive power holds if used by another technology.

## Results

### e-DNA profiling

The closed-reference assignment of reads resulted in four different readcount matrices for each sample. In addition, we also applied the more conventional open-reference approach to the Illumina data using three different methods (Fig. 1), resulting in a total of seven read count matrices per sample. Summary statistics for readcount matrices can be found in Table 1.

**Table 1:** The table summarizes the read count matrices generated using different approaches. The Tool refers to bioinformatic softwares used to assign/group reads. Mean readcount is the average number of reads per sample.

| Technology | Tool | Database | Mean readcount | Number of features |
| --- | --- | --- | --- | --- |
| Illumina | EMU | AQUAeD-DB | 52414 | 4509 |
| Nanopore | EMU | AQUAeD-DB | 54685 | 2832 |
| Illumina | VSEARCH | AQUAeD-DB | 46063 | 10939 |
| Nanopore | VSEARCH | AQUAeD-DB | 31404 | 5741 |
| Illumina | Dada2 | - | 35291 | 18604 |
| Illumina | OTU97 | - | 46743 | 48524 |
| Illumina | UNOISE | - | 46812 | 92808 |

### Effects on prediction errors

Comparison of absolute nEQR prediction error distributions for the various combinations of sequencing technologies, read assignment or clustering tools and machine learning approaches demonstrated that the difference between Illumina and Nanopore technology is minimal regardless of the tool used for read assignment or the ML method (Fig. 2).

**Figure 2:**
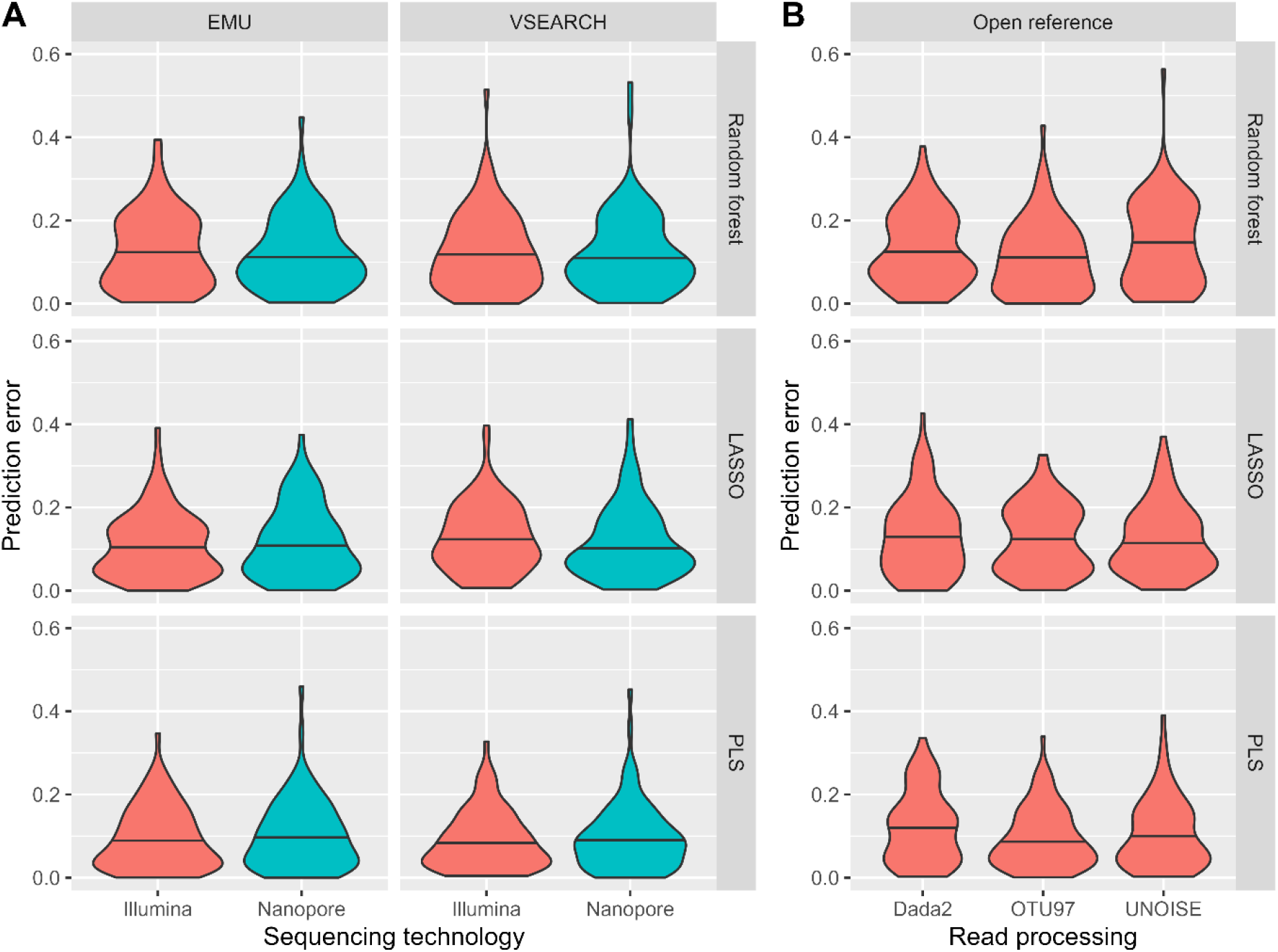
The violin plots demonstrate the prediction error for closed-reference (panel A) and open-reference analysis (panel B) for three different ML algorithms. The horizontal line in each violin indicates the median value of prediction error. The x-axis in panel A shows the sequencing technology used for EMU and VSEARCH. The x-axis in panel B shows the read processing approach used.

The closed-reference analysis yielded comparable prediction errors for both the Random Forest (RF, error = mean ± 0.13) and LASSO (error = mean ± 0.12) models (Fig. 2, panel A). In contrast, the Partial Least Squares Regression (PLSR) consistently outperformed the other machine learning algorithms for both the Illumina and Nanopore datasets, as reflected by a significantly lower mean prediction error (error = mean ± 0.10). An ANOVA on prediction errors as the response variable revealed no significant effects from sequencing technology (Illumina or Nanopore) or read assignment tool (EMU or VSEARCH). There was a significant effect from the ML approach, with PLS (p<0.05) outperforming Random Forest and LASSO, but the differences were small to be practically relevant.

For open-reference analysis, prediction error results were similar for all three methods tested (OTU97, UNOIOSE, and Dada2; error = mean ± ∼0.1) (Fig. 2, panel B). Open-reference analysis did not result lower prediction errors as compared to closed-method analysis, despite the identification of larger numbers of features by open-reference methods (Table 1).

### Feature selection and feature assessment

Given the small but significant differences in nEQR prediction errors, subsequent analysis focused on improving predictions for both Illumina and Nanopore data by applying feature selection using LASSO with jackknife. Feature selection resulted in identification of 36 taxa from Illumina data and 43 taxa from Nanopore data, of which four taxa were shared between feature sets. The positive and negative regression coefficients from the PLS algorithm, was evenly distributed across the selected features (Fig. 3).

**Figure 3:**
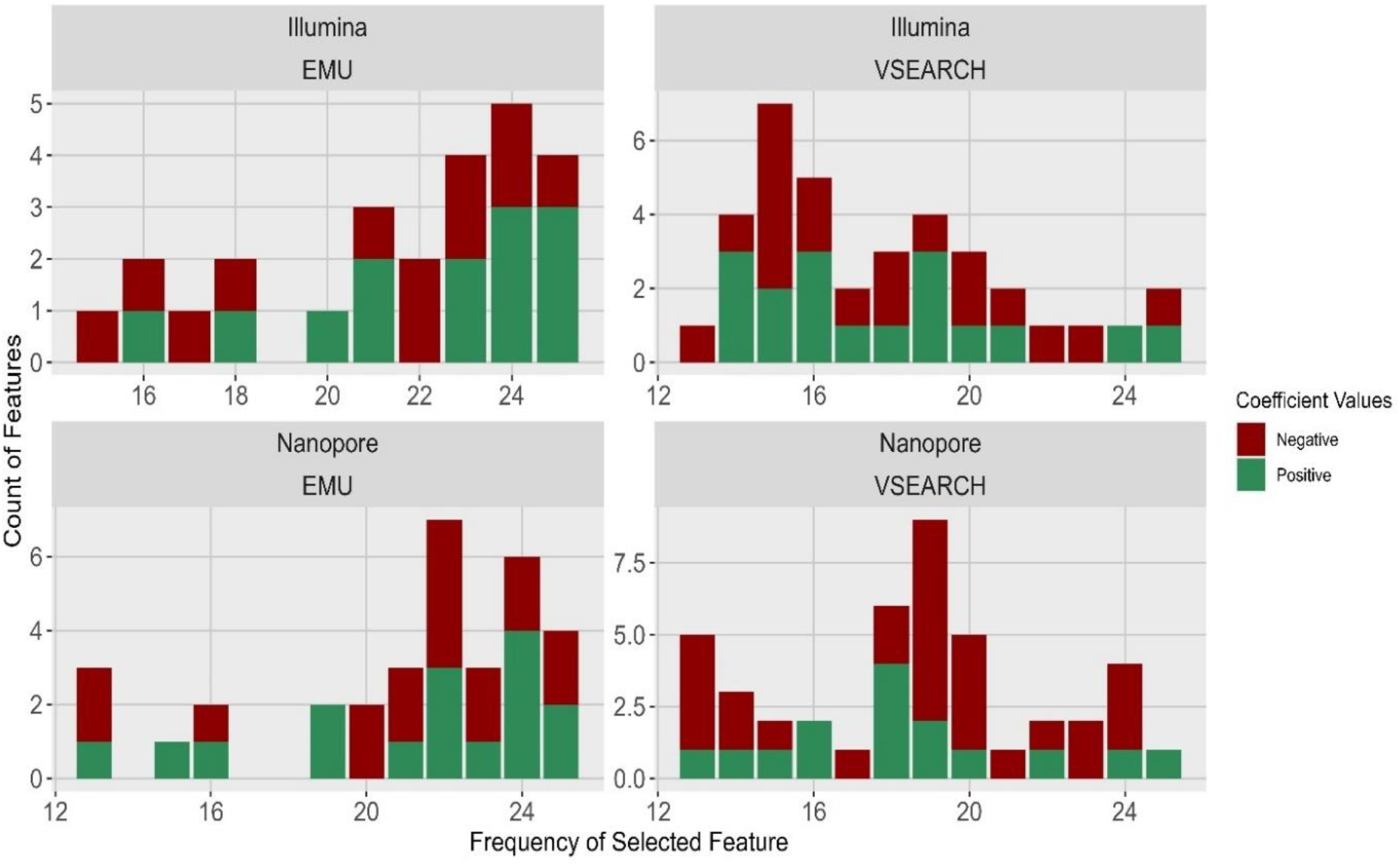
The bar chart displays the number of features selected from 13 to 25 times (x-axis) for both Illumina and Nanopore using EMU and VSEARCH. Colours represent the positive and negative regression coefficient values for each feature, i.e. if they have a positive or negative correlation to ecosystem status.

### Improved prediction

From this feature selection we extracted the relevant Nanopore and Illumina sequence reads for assignment using both EMU and VSEARCH. Using these reduced feature matrices, a PLSR model was fitted for each matrix, applying the same CLR transformation and cross-validation procedure as previously described. The prediction error from PLSR on the optimized features for Illumina sequences reveals that predictions exhibit a minimal mean error of ± 0.04. The predictions on Nanopore data shows an approximate mean error of ± 0.05. The predictions generated from the reduced feature matrix using the PLSR algorithm for both Nanopore and Illumina data (see Figure 4) yield the best results. Again, the differences between Illumina and Nanopore are very small compared to method variations for each sequencing technology (Fig. 5).

**Figure 4:**
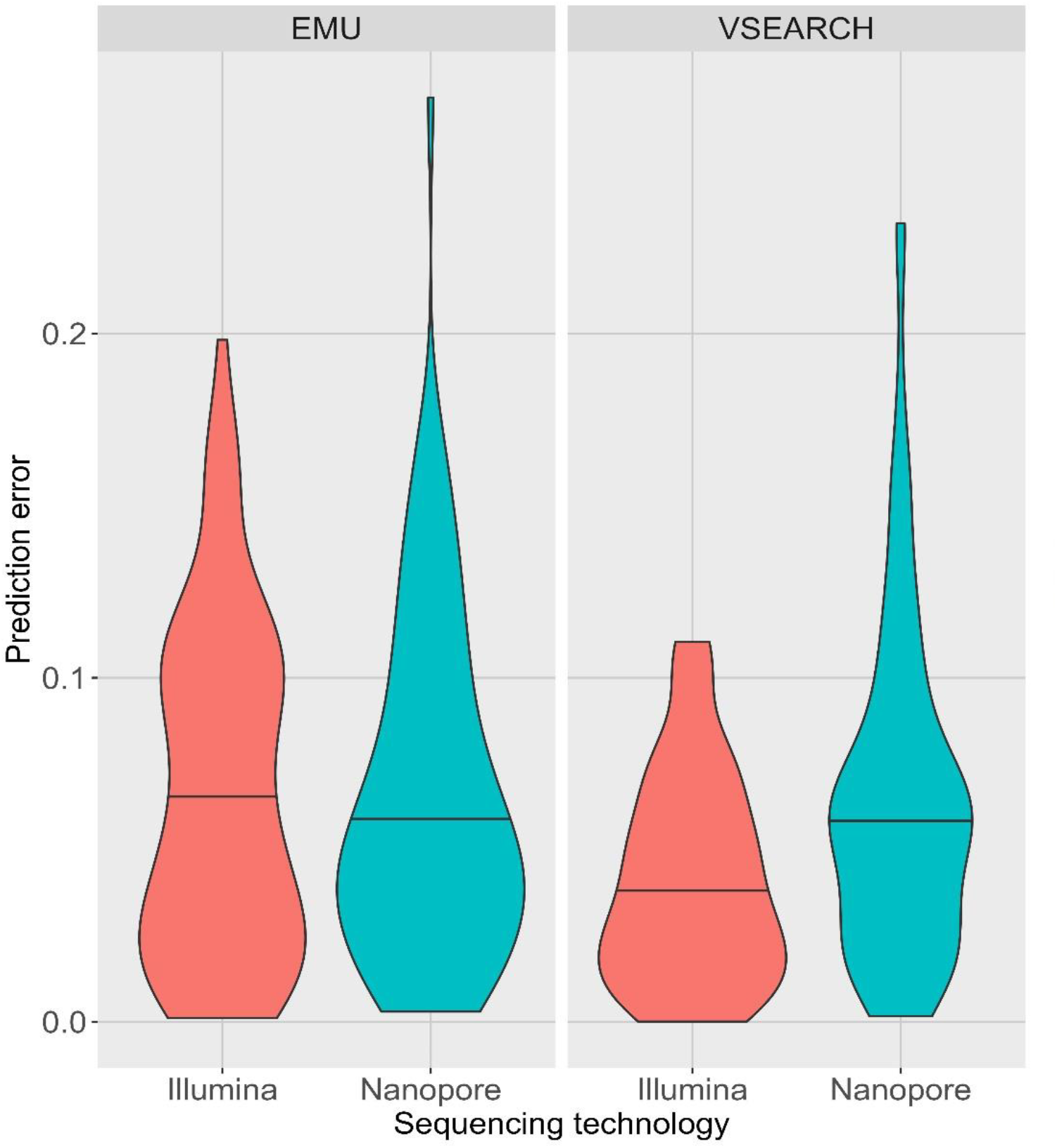
Prediction error distributions. Violin plots demonstrate the prediction error for Illumina (red) and Nanopore (blue) analysis when using EMU (left panel) or VSEARCH (right panel) for assignment. The horizontal line in each violin indicates the median value.

**Figure 5:**
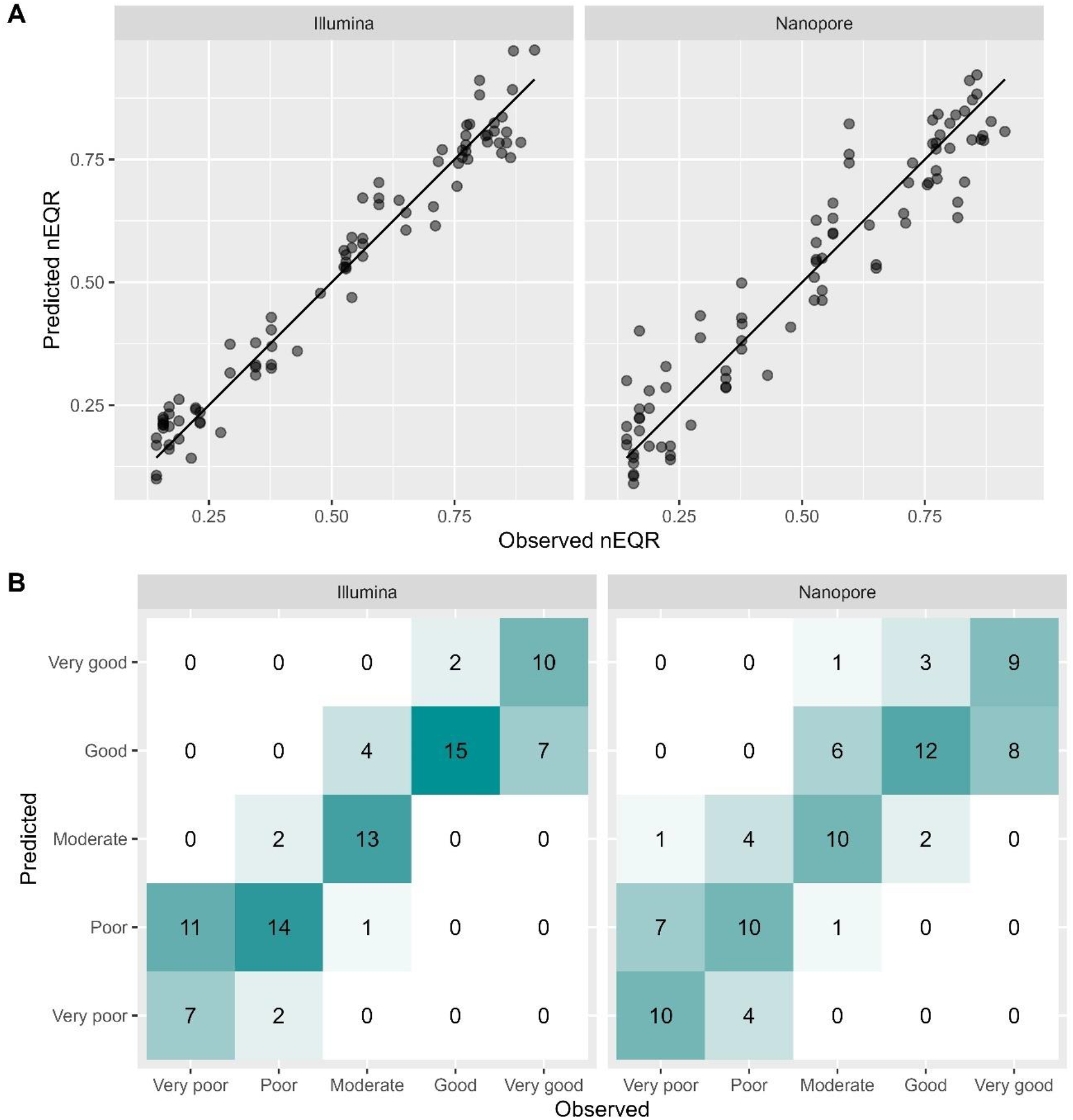
Observed versus predicted nEQR values. (A) Scatterplots showing observed versus PLS-predicted nEQR values for VSEARCH-assigned Illumina (left panel) and Nanopore (right panel) reads for each sample. The Illumina data exhibits a correlation of 0.98, while the Nanopore data demonstrates a correlation of 0.95 with VSEARCH. Panel B shows the observed versus PLS-predicted nEQR values for the same data in A, divided by nEQR status classification. The value in each cell is the number of samples falling into each combination of predicted-observed category.

The selected features from Illumina and Nanopore data exhibited only minimal overlap (four taxa). Given the high correlation among many features in these two datasets, however, the selected features may demonstrate similar predictive power. To investigate this, we conducted an analysis by swapping the selected features between the Illumina and Nanopore. When the features were applied to their respective datasets, the correlations between observed and predicted values were as follows: VSEARCH produced a correlation of 0.98 for Illumina (mean prediction error: ±0.04) and 0.95 for Nanopore (mean prediction error: ±0.06), while the EMU yielded correlations of 0.95 for Illumina (mean prediction error: ±0.06) and 0.95 for Nanopore (mean prediction error: ±0.06). When the feature sets were swapped, the corresponding correlations were 0.92 for Illumina (mean prediction error: ±0.07) and 0.88 for Nanopore (mean prediction error: ±0.09) with VSEARCH, and 0.90 for Illumina (mean prediction error: ±0.08) and 0.87 for Nanopore (mean prediction error: ±0.10) with EMU.

## Discussion

This study has demonstrated the potential for accurate nEQR prediction based on 16S rRNA Nanopore sequencing of seafloor sediments The predictions and the ANOVA results from different combinations of assignment and machine learning data analysis approaches indicate that the ecosystem status predictions based on Nanopore data are comparable to those based on Illumina data (Figure 2). Longer sequence reads, such as those obtained using Nanopore technology, can potentially improve taxonomic resolution of sequencing-based approaches (Benítez-Páez et al., 2016; Krehenwinkel et al., 2019; Nygaard et al., 2020). In combination with continual technological developments to decrease sequencing error rates, Nanopore may emerge as an acceptable choice for future amplicon sequencing-based environmental monitoring initiatives (Urban et al., 2021).

One notable finding is that the two different tools used to assign sequence reads to reference database sequences produced similar results (Figure 2) despite their specialized designs for specific technologies. VSEARCH (Rognes et al., 2016) has been designed for analysis of short-read Illumina data while EMU (Curry et al., 2022) was developed for analysis of long-read Nanopore. The VSEARCH tool generates larger number of features (Table 1), while the EMU tool produces fewer features with higher read count. Nevertheless, results of this study indicate that both tools perform comparably well for both Illumina and Nanopore sequence data. One possible explanation for the comparable predictive value of Nanopore and Illumina data is recent improvements made to Nanopore sequencing accuracy (Zhang et al., 2024). This may explain why VSEARCH, despite being specifically designed for Illumina, works well with Nanopore data, whereas EMU, developed a few years earlier, may not have been optimized for the improved accuracy now seen in Nanopore reads.

Both open- and closed-reference assignment workflows were used for the analysis of Illumina data, despite the fact that open-reference approaches without pre-defined reference sequences are more commonly employed (Terrat et al., 2020). nEQR error prediction using machine learning and ANOVA clearly indicated, however, that open-reference denoising and clustering had no significant impact on prediction accuracy when compared to closed-reference approaches (Figure 2). The three commonly-used open-reference methods DADA2, UNOISE and 97% identity clustering, identified a larger number of features (Table 1), however the predictive power with respect to ecosystem status was statistically indistinguishable from closed-reference approaches. Moreover, to standardize surveillance, a closed-reference approach seems essential in any circumstance. To ensure comparability of taxonomic profiles over time and locations, a fixed set of references are required.

In general, feature selection with machine learning was effective in reducing the dimensionality readcount matrices. The number of selected features for Illumina and Nanopore data varies, yet some common features are observed. Some differences are expected due to the use of different primers for each technology (Abellan-Schneyder et al., 2021), but more likely the differences are due to unstable selection (Papoutsoglou et al., 2023). Even though the specific taxa vary between datasets, they frequently represent similar types of organisms. This is consistent with data where many features are correlated, and only one representative is selected from each group of correlated features (Dommann et al.). While the specific representative may change, it still reflects the same group. In these instances, even minor changes in the data lead to variability in the feature selection (Hernández Medina et al., 2022). However, the selected features show consistent distribution of positive and negative regression coefficients (Figure 3) with a distinct majority of the negative coefficients. This suggests that even if the selected features vary, they seem to represent a similar set of taxa with positive/negative association to environmental impact.

The comparison of correlations between observed and predicted nEQR values for both sequencing technologies, using selected features from their respective datasets versus swapping features across platforms, indicates a slight drop in correlation, particularly for Nanopore (Fig. 5). This outcome is expected, as the features utilized are no longer optimal. Nonetheless, the PLSR predictions demonstrate an effective application of these features, resulting in improved accuracy compared to using all variables. The feature selection reduced the number of features for both Illumina and Nanopore. Comparison of correlation results reveal that these features are more similar after all, they convey similar information despite being different features. Some drop in predictive power is expected due to the fundamental differences between Illumina and Nanopore data, as indicated in previous publication (Philip et al., 2024). All in all, this essentially means that if ecosystem monitoring is to be done by Nanopore sequencing, features should be selected using closed-reference methods with a Nanopore reference database to obtain optimal results.

The feature selection using LASSO regression improved the predictions as evident in the results from PLS fitting on the reduced feature matrix (Figure 4). Prediction errors have been significantly decreased from approximately ±0.10 to ±0.04 after the feature selection. Both the Illumina and Nanopore approaches have robust nEQR prediction capabilities, as shown in Figure 5, which presents their optimised performance using VSEARCH. The data points in panel A of Figure 5 closely align with the diagonal line, indicating a strong positive correlation between predicted and observed values. The confusion matrix in panel B also present the performance of both Illumina and Nanopore.

Given the labour-intensive manual sorting and taxonomic processes involved in obtaining nEQR values using morphotaxonomic analysis, it is important to acknowledge potential uncertainty and bias associated with traditional taxonomic approaches (Duarte et al., 2023; Hering et al., 2018). It would therefore be optimistic to assume that any other method, for example sequencing, can perfectly predict nEQR (Cordier et al., 2017; Cowart et al., 2015; Ruppert et al., 2019). There is currently little data to support uncertainty estimates for nEQR calculations (Clinton et al., 2024). The feature selection results in this study are promising, however features were selected from the same 88 samples used for prediction, suggesting that the features identified are ‘ideal’ for this dataset. Although this illustrates the diagnostic potential of an appropriate feature set, universal application would require feature selection based on a significantly larger dataset.

Previous research has proven the capabilities of Illumina-based prediction studies, emphasizing their effectiveness in applications related to seafloor health prediction (Cordier et al., 2017). These findings suggest that, similar to Illumina, Nanopore is also a reliable option for predictive approaches of this nature. Furthermore, the results show that the resolutions currently generated from Nanopore data enable the models to elucidate robust connections between microbial profiles and ecological status.

## Conclusion

This work aimed to evaluate the predictive capabilities of microbial communities based on both Nanopore and Illumina sequencing platforms in determining the ecological status of seafloor sediment samples. This study indicates that Nanopore-based microbial community profiles are comparable to Illumina with regard to reliability for predictive applications. Furthermore, there are indications that with advancements in technology, Nanopore sequencing has the potential to outperform Illumina when working with larger and more diverse seafloor sample datasets. In addition, the implementation of feature selection is essential for establishing a standardized approach and for generating best fitted models with minimized prediction errors. Ongoing advancements in Nanopore technology has the potential to provide higher resolution amplicon data for cost-effective ecological assessments and monitoring.

## Acknowledgements

We extend our gratitude to the fish farmers who supplied samples for our study. We would also like to thank the STIM, Aqua Kompetanse, and Akvaplan-NIVA employees that participated in the samples collection.

## Data availability

All raw sequences used in this study have been deposited in the NCBI BioProject database. Illumina 16S rRNA sequences is a part of the project with the accession code PRJNA1128851. The raw Nanopore sequencing data for subset B has been uploaded with the accession code [PRJNA1153974](https://www.ncbi.nlm.nih.gov/bioproject/1153974). The AQUAED-DB and taxonomy details are publicly available (https://arken.nmbu.no/~pmelcy/share/AQUAED-DB/). The metadata files for all illumina and Nanopore processing are available in https://arken.nmbu.no/~pmelcy/share/AQUAED-DB/metadata/metadata_illumina_subsetB.txt and https://arken.nmbu.no/~pmelcy/share/AQUAED-DB/metadata/metadata_nanopore_subsetB.txt. The metadata file and readcount matrices for predictions are available in https://arken.nmbu.no/~pmelcy/share/Predictions/.

## Author contributions

Knut Rudi, Lars Snipen and Melcy Philip conceived the study; Jessica Louise Ray, Nigel Keeley, Morten Stokkan, Ragnhild Pettersen and Sanna Majaneva design the collection of sediment samples; Tonje Nilsen performed all lab experiments; Lars Snipen and Melcy Philip performed all analyses, with conceptual input from Knut Rudi; Knut Rudi, Lars Snipen and Melcy Philip contributed to interpretation of results; Knut Rudi, Lars Snipen and Melcy Philip drafted the manuscript. All author(s) read and commented on the manuscript.

## Funding

This study is a part of the project: “AQUAeD”, funded by The Research Council of Norway (Project Number: 320076).

## Competing interests

At the time of manuscript writing, Sanna Majaneva, Ragnhild Pettersen were employed by company Akvaplan-niva AS. Jessica Louise Ray was employed by company Aqua Kompetanse AS and Morten Stokkan was employed by company STIM AS. The remaining authors declare that the research was conducted in the absence of any commercial or financial relationships that could be construed as a potential conflict of interest.

## Notes

https://arken.nmbu.no/~pmelcy/share/AQUAED-DB/

https://arken.nmbu.no/%7Epmelcy/share/Predictions/

